# Insight into the evolutionary assemblage of cranial kinesis from a Cretaceous bird

**DOI:** 10.1101/2022.07.13.499923

**Authors:** Min Wang, Thomas A. Stidham, Jingmai K. O’Connor, Zhonghe Zhou

## Abstract

The independent movements and flexibility of various parts of the skull, called cranial kinesis, is an evolutionary innovation that is found in living vertebrates only in some squamates and crown birds, and considered to be a major factor underpinning much of the enormous phenotypic and ecological diversity of living birds, the most diverse group of extant amniotes. Compared to the postcranium, our understanding of the evolutionary assemblage of the characteristic modern bird skull has been hampered by sparse fossil records of early cranial materials, with competing hypotheses regarding the evolutionary development of cranial kinesis among early members of the avialans. Here, a detailed three-dimensional reconstruction of the skull of the Early Cretaceous enantiornithine *Yuanchuavis kompsosoura* allows for its in depth description, including elements that are poorly known among early diverging avialans but are central to deciphering the mosaic assembly of features required for modern avian cranial kinesis. Our reconstruction of the skull shows evolutionary and functional conservation of the temporal and palatal regions by retaining the ancestral theropod dinosaurian configuration within the skull of this otherwise derived and volant bird. Geometric morphometric analysis of the palatine suggests that loss of the jugal process represents the first step in the structural modifications of this element leading to the kinetic crown bird condition. The mixture of plesiomorphic temporal and palatal structures together with a derived avialan rostrum and postcranial skeleton encapsulated in *Yuanchuavis* manifests the key role of evolutionary mosaicism and experimentation in early bird diversification.

## Introduction

The growing fossil record of Mesozoic avialans has revealed that the initial appearance of many evolutionary novelties associated with living birds in fact originated among early lineages (*Xu et al., 2014; Brusatte et al., 2015; Chiappe and Meng, 2016*). Much of the research interest in the early evolution of birds has focused on flight relevant morphologies (*Sullivan et al., 2017; Heers et al., 2021*). However, little is known about how the bulky and akinetic dinosaurian skull became transformed into the delicate, kinetic, and functionally diverse extant bird skull (*Bhullar et al., 2016*), which is inferred to have contributed significantly to their spectacular adaptive radiation, culminating in the >10,000 species recognized today. This lack of resolution results in part from the rarity of cranial materials of early avialans and the limitd data available from conventional methods of examination given their typically poor and two-dimensional preservation (*Witmer and Martin, 1987; O’Connor and Chiappe, 2011*). Meanwhile, some previous studies have perhaps overzealously overstated the cranial resemblance between stem and crown birds, including positing the existence of certain bony contacts and intracranial joints required for kinesis, though yet to be confirmed positively in fossil taxa (*Witmer and Martin, 1987; Holliday and Witmer, 2008; Wang et al., 2021a*). As a step forward in our understanding of avialan cranial evolution, we applied x-ray computed tomography scanning of the recently described Early Cretaceous bird *Yuanchuavis kompsosoura* (*Supplementary videos 1 and 2*) (*Wang et al., 2021b*). *Yuanchuavis* phylogenetically is assigned to the second earliest branching enantiornithine clade (Pengornithidae), and preserves a unique pintail feather morphotype that has been inferred to be used for sexual display (*Wang et al., 2021b*). Our study documents previously unrecorded and unrecognized cranial features with clear functional and macroevolutionary implications.

## Results

Like other pengornithids (*Zhou et al., 2008; Hu et al., 2015; O’Connor et al., 2016*), the premaxillae are unfused. As in most Early Cretaceous enantiornithines including other pengornithids except for *Chiappeavis* (*Zhou et al., 2008; Hu et al., 2015; O’Connor et al., 2016*), the short premaxillary corpus is rostrocaudally longer than dorsoventrally high, with subparallel dorsal and ventral margins (*Figures 1, 2a; Figure 1—figure Supplement 1*). The maxillary process of the premaxilla extends beyond the rostrocaudal midpoint of the external naris, which is proportionately longer than the state in other pengornithids where it terminates caudally within the rostral fourth of the external naris (*Zhou et al., 2008; Hu et al., 2015; O’Connor et al., 2016*). Six premaxillary teeth are present on each side (*Figure 2b*), not five as estimated in a previous study (*Wang et al., 2021b*), and this is greater than the maximum number of four in other avialans (*Louchart and Viriot, 2011; O’Connor and Chiappe, 2011; Hu et al., 2019*). As in other pengornithids (*Zhou et al., 2008; Hu et al., 2015; O’Connor et al., 2016*), the tooth crowns are weakly expanded and the tapered apices are slightly recurved. The frontal processes of the premaxillae become dorsoventrally compressed as they extend caudally, and the distal quarter of the process is caudolaterally tapered such that the two premaxillae define a medial notch for the nasals. The triradiate maxilla constitutes the major portion of the facial margin as in other early avialans (*Figure 2c*) (*O’Connor and Chiappe, 2011; Wang et al., 2021a*). The ascending process lacks the fenestra present in *Pengornis* (*Zhou et al., 2008*). The broad jugal process constricts sharply along its caudal fourth, a unique feature otherwise unknown among early diverging avialans (*O’Connor and Chiappe, 2011; Rauhut et al., 2018; Kundrát et al., 2019; O’Connor et al., 2020; Wang et al., 2021a*), although it is likely that this portion of the maxilla is commonly broken and lost or covered by the jugal. The ventral margin of the jugal process flares laterally, forming a lateral groove that accommodates the jugal. Nine maxillary teeth are present (seven preserved teeth and two additional alveoli preserved in the right maxilla), which like other pengornithids exceeds the number present in some enantiornithines (e.g., bohaiornithids) (*O’Connor and Chiappe, 2011; Wang et al., 2014*). The exact number of maxillary teeth is unclear in other pengornithids.

**Figure 1.**
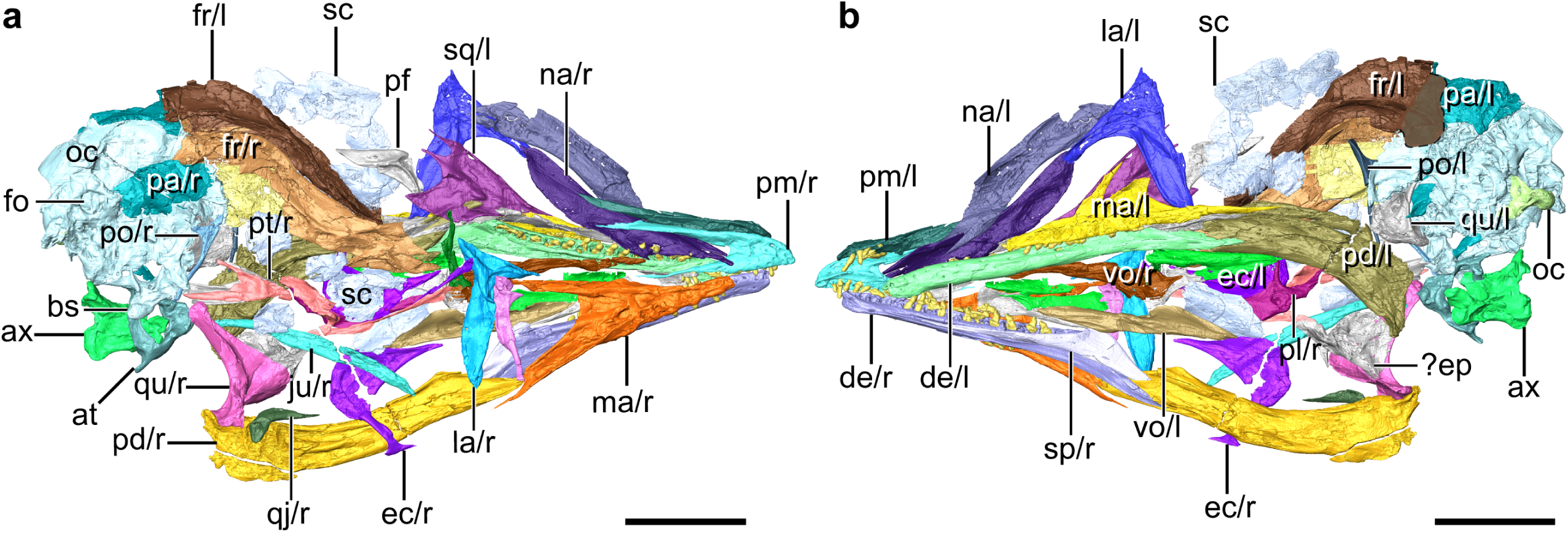
Digital reconstruction of the skull of *Yuanchuavis*, IVPP V27883. **a, b,** Skull in right (**a**) and left (**b**) view, respectively. ax, axis; at, atlas; bs, basipterygoid process; de, dentary; ec, ectopterygoid; ep, epipterygoids; fo, foramen magnum; fr, frontal; ju, jugal; la, lacrimal; ma, maxilla; na, nasal; oc, occipital region; op, occipital condyle; pa, parietal; pd, post-dentary mandible; pl, palatine; pm, premaxilla; po, postorbital; pt, pterygoid; qj, quadratojugal; qu, quadrate; sc, scleral ossicles; sp, splenial; sq, squamosal; vo, vomer; and r/l, right/left side. Scale bars, 10 mm. **Figure supplementary 1.** Cranial anatomy of *Yuanchuavis*, IVPP27883. **Figure supplementary 2.** Digital reconstruction of atlas and axis of *Yuanchuavis*.

**Figure 2.**
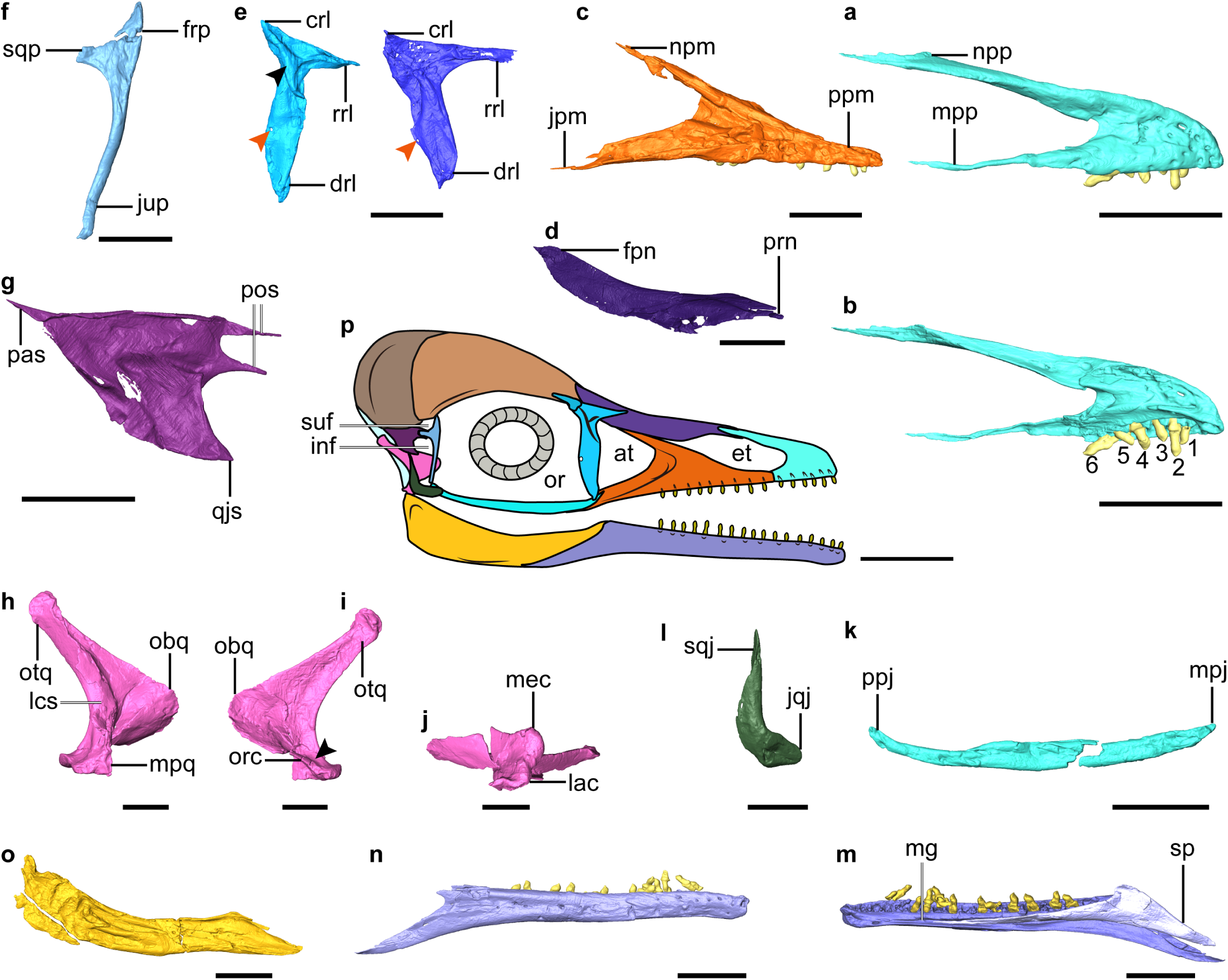
Digital reconstruction of facial and jaw bones of *Yuanchuavis*. **a, b,** Right premaxilla (numbers 1**–**6 denote premaxillary teeth). **c,** Right maxilla. **d,** Right nasal. **e,** Right (right column) and left lacrimal in lateral and medial view, respectively (black and red arrowhead denote the lateral flange and foramen, respectively). **f,** Right postorbital. **g,** Left squamosal. **h–j**, Right quadrate (**i;** arrowhead denotes the absence of a pterygoid condyle). **k,** Right jugal. **l**, Right quadratojugal. **m, n,** Right dentary. **o,** Right post-dentary mandible. **p,** Reconstruction of *Yuanchuavis* skull in lateral aspect. **a**, **c–f**, **h**, **k–m**, **o**: Lateral view; **b, g, i, n**: medial view; **j**: ventral view. at, antorbital fenestra; crl, caudal ramus of lacrimal; drl, descending ramus of lacrimal; et, external naris; frp, frontal process of postorbital; inf, infratemporal fenestra; jpm, jugal process of maxilla; jqj, jugal process of quadratojugal; jup, jugal process of postorbital; lac, lateral condyle; lsc, lateral crest; mec, medial condyle; mg, Meckel’s groove; mpj, maxillary process of jugal; mpp, maxillary process of premaxilla; mpq, mandiblar process of quadrate; npm, nasal process of maxilla; npp, nasal process of premaxilla; obq, orbital process of quadrate; or, orbit; orc, orbitocondylar crest; otq, otic process of quadrate; pas, paraoccipital process of squamosal; pf, prefrontal; pos, postorbital process of squamosal; ppj, postorbital process of jugal; ppm, premaxillary process of maxilla; qjs, quadratojugal process of squamosal; rrl, rostral ramus of lacrimal; sp, splenial; sqj, squamosal process of quadratojugal; sqp, squamosal process of postorbital; suf, supratemporal fenestra. Scale bars, 5 mm (**a–e, g–k, m–o**), 2.5 mm (**f, l**), 10 mm (**p**). **Figure supplementary 1.** Additional cranial anatomy of *Yuanchuavis*.

The nasal is weakly bowed ventrally and tapered at both ends (*Figure 2d*). A maxillary process, present in *Parapengornis* and other enantiornithines (*Zhou et al., 2005; O’Connor and Chiappe, 2011; Li et al., 2021*), is absent. The elongate nasals contact medially for almost their entire length such that the premaxillae and frontals do not contact. The rostral end of the nasal is forked as in *Pengornis* (*Zhou et al., 2008*), and receives the caudal tapered portion of the nasal process of the premaxilla (*Figure 2—figure Supplement 2a, b*). The nasal has a width to length ratio of 0.11, notably more slender than in some enantiornithines (e.g., *Eoenantiornis* and *Protopteryx*), but wider than in *Falcatakely* (*O’Connor et al., 2020*). In contrast to *Pengornis* (*Zhou et al., 2008*), the bone is not perforated by a foramen.

The lacrimal is triradiate with a robust descending ramus that is longer and wider than the more delicate rostral ramus, which in turn is twice the length of the caudal ramus (*Figure 2e*). The straight dorsal margin of the lacrimal lacks the central concavity characteristic of *Pengornis* (*Zhou et al., 2008*). The descending ramus widens rostrocaudally along its central portion, and then narrows towards the ventral end, lacking the ventral expansion seen in *Falcatakely* and *Ichthyornis* (*O’Connor et al., 2020; Torres et al., 2021*). An oval fenestra is developed near its caudal margin at the dorsoventral midpoint (recognized in both sides), which is absent in other early avialans including other pengornithids (*Mayr et al., 2005; O’Connor and Chiappe, 2011; Zhou et al., 2013; O’Connor et al., 2020*). The lacrimal bears a triradiate lateral embossment forming crests along each ramus, and it is centered at the juncture of the three rami. No similar structure hitherto has been described among other early avialans (*O’Connor and Chiappe, 2011; Rauhut et al., 2018; Wang et al., 2021a*), but this morphology is present in deinonychosaurians (*Norell et al., 2006; Xu et al., 2015*). While the dorsal branches of the lateral embossment are relatively straight, the ventral branch curves with a concave rostral margin, and it decreases in height towards its tip, ending in the dorsal one-fifth of the rostral margin of the descending ramus. The preserved left prefrontal is similar to the condition in *Archaeopteryx* (*Rauhut et al., 2018*) and consistent with the relatively short proportion of the caudal ramus of the lacrimal (*Figure 1*).

The postorbital is an enigmatic element poorly known among enantiornithines (likely independently reduced in some taxa (*Xu et al., 2021*), but is preserved exquisitely here (*Figure 2f*). The Y-shaped postorbital has a pointed frontal process and a blunter squamosal process that are oriented at a near right angle to one another. The dorsal margin between the frontal and squamosal processes is concave, forming the rostroventral margin of the supratemporal fenestra. The squamosal process is shorter than the frontal process and ends in a blunt articular surface for the squamosal. The jugal process is approximately four times longer than the frontal process, delicate, and deflected caudoventrally. The squamosal is another poorly known cranial element in enantiornithines that has been recognized in only two specimens (LP-4450-IEI, and IVPP V12707) (*O’Connor and Chiappe, 2011*). Compared to those specimens, the squamosal of *Yuanchuavis* is morphologically more similar to that of *Archaeopteryx* and non-avialan theropods such as *Deinonychus*, attesting to its relatively large size, the robust quadratojugal process, and the rostrally deeply forked postorbital process (*Figure 2g*). In contrast, the postorbital process is not forked in LP-4450-IEI and IVPP V12707 (*O’Connor and Chiappe, 2011; Wang et al., 2021a*). The notch in the postorbital process is deeper than in *Archaeopteryx* (*Elzanowski and Wellnhofer, 1996*), but not as pronounced as in *Deinonychus* (*Ostrom, 1969*). As in *Archaeopteryx* (*Elzanowski and Wellnhofer, 1996*), the dorsal and ventral branches of the postorbital process are subequal in length; whereas the ventral branch is longer in *Deinonychus* (*Ostrom, 1969*). The trapezoidal shaped quadratojugal process bears a pointed rostroventral corner that projects as far rostrally as the postorbital process. In contrast, the quadratojugal process is slender and rod-like in *Archaeopteryx* and other enantiornithines, and is triangular in non-avialan theropods (*Ostrom, 1969; Norell et al., 2006; O’Connor and Chiappe, 2011; Xu et al., 2015; Wang et al., 2021a*). The paraoccipital process is sharply tapered as it projects caudally, and it is connected with the quadratojugal process by a roughly straight ventral margin, resembling the condition in *Linheraptor* (*Xu et al., 2015*). By contrast, that process is caudoventrally directed, resulting in a right angle with the quadratojugal process in IVPP V12707 and *Deinonychus* (*Ostrom, 1969; Wang et al., 2021a*).

The unfused frontals and parietals are crushed, revealing few features except that the skull roof is vaulted dorsally. There is an elongate ridge along the lateral edge of the frontal forming a concave margin around the dorsal margin of the orbit, and this ridge runs for most of the length of the frontal. The frontal does not appear to have contributed to a postorbital process.

As in other early avialans and non-avialan theropods (*Turner et al., 2012; Hendrickx et al., 2015; Wang et al., 2021a*), the quadrate has a bicondylar mandibular process (*Figure 2h–j*, *Figure 2—figure supplement* 2c–e). Like some enantiornithines but not *Longipteryx* (*O’Connor and Chiappe, 2011; Stidham and O’Connor, 2021*), the medial condyle is larger than the lateral one (*Figure 2j*). The caudal surface of the quadrate corpus is not perforated by a foramen as in some enantiornithines including *Pengornis* and *Shenqiornis* (*Zhou et al., 2008; O’Connor and Chiappe, 2011*). In lateral view, the caudal margin of the quadrate is concave and bowed with the medial and lateral mandibular condyles positioned caudally relative to the corpus as in the juvenile enantiornithine IVPP V12707 (*Wang et al., 2021a*), rather than being level with the shaft as in non-avialan theropods such as *Linheraptor* (*Turner et al., 2012; Xu et al., 2015*). The bowing is restricted to the ventral half of the quadrate with the caudal margin (lateral view) being relatively straight in the dorsal half, and the absence of cracks in the corpus supports this as the original morphology (*Figure 2h, j*). This morphology also suggests that the quadrate was inclined, with the otic process positioned caudal to the mandibular process. There is a shallow concavity separating the medial and lateral condyles near the mediolateral midpoint of the mandibular process (*Figure 2—figure Supplement 1c*). The lateral edge of the mandibular process is broken so that the morphology of the articulation with the quadratojugal cannot be identified in detail, but the remnants are directed and extended somewhat rostrally, in a manner similar to the state present in *Linheraptor* where it has rostrocaudally elongate contact with the quadratojugal. That elongation has not been identified in other enantiornithines. This rostrally directed process is ventral to the ventral edge of the orbital process. The caudal surface of the quadrate corpus forms a sharp crest along its dorsoventral length from the otic head ventrally to just dorsal to the concavity between the medial and lateral mandibular condyles. That sharp caudal crest is present in many, but not all enantiornithines. Like IVPP V12707 and non-avialan theropods (*Norell et al., 2006; Hendrickx et al., 2015; Wang et al., 2021a*), a distinct pterygoid condyle is absent on the quadrate (*Figure 2—figure Supplement 2c*), indicating the absence of a condylar-based joint between the quadrate and pterygoid, a derived feature present in *Ichthyornis*, hesperornithiforms, and crown taxa (*Gingerich, 1976; Baumel and Witmer, 1993; Field et al., 2018a*). The shape of the orbital process is nearly identical to that of *Archaeopteryx* and non-avialan theropods (called the pterygoid ramus) in having a rounded and blunt convex rostral apex with a straight dorsal margin that extends to the otic process (*Figure 2h*) (*Currie, 1995; Norell et al., 2006; Hendrickx et al., 2015; Rauhut et al., 2018*). In contrast, the orbital process is dorsoventrally thin and tapers rostrally in other enantiornithines (*Sander et al., 2001; Stidham and O’Connor, 2021; Wang et al., 2021a*), which is further modified into a narrower, pointed process in *Ichthyornis* and more crownward taxa (*Elzanowski and Stidham, 2011; Field et al., 2018a; Torres et al., 2021*). As in other enantiornithines and most non-avialans (*Stidham and O’Connor, 2021; Wang et al., 2021a*), the otic head is not divided into separate squamosal and otic capitula. The otic process has a triangular cross section, with the caudal crest on the corpus flanked by flattened medial and lateral surfaces, as in other enantiornithines.

A fragment preserved and compressed to the medial surface of the orbital process of the right quadrate is tentatively identified as the epipterygoid (*Figure 1*), given its proximity with the right pterygoid and current overlapping state with the quadrate. No epipterygoid has been reported previously among early diverging avialans, but it is known in some avialan outgroups.

As in *Pterygornis* (*Wang and Hu, 2017*), the caudal end of the jugal curves caudodorsally rather than being forked as in other pengornithids and enantiornithines such as *Falcatakely* and IVPP V12707 (*Figure 2k*) (*O’Connor and Chiappe, 2011; Wang et al., 2021a*). The L-shaped quadratojugal has a dorsally directed postorbital process (with a groove on its caudal dorsal edge presumably for articulation with the squamosal) and a short jugal process (*Figure 2l*).

The pterygoid has a large, forked, and caudodorsally directed quadrate ramus (*Figure 3a, b*), and the pterygoid does not contact the quadrate on its ventromedial aspect through a condyle as in ornithurines (*Baumel and Witmer, 1993*). The occurrence of this morphology in *Yuanchuavis* reinforces the recent identification of the retention of this plesiomorphic curved and forked non-avialan pterygoid shape among enantiornithines (*Figure 3—figure Supplement 1*) (*Wang et al., 2021a*). The mediolaterally compressed quadrate ramus is forked into a longer dorsal and shorter ventral processes, opposite of the condition seen in the enantiornithine IVPP V12707 and dromaeosaurids (*Figure 3— figure supplement 1a–d*) (*Ostrom, 1969; Barsbold and Osmólska, 1999; Tsuihiji et al., 2014; Wang et al., 2021a*). By contrast, the ornithurine pterygoid is modified substantially with a reduced or absent quadrate ramus and an overall elongation reaching to the area just dorsal to the medial mandibular condyle of the quadrate, as seen in the triangular shape in hesperornithiforms and overall more strut-like appearance in crown birds (*Figure 3—figure Supplement 1e*) (*Gingerich, 1976; Baumel and Witmer, 1993; Zusi and Livezey, 2006*). A potentially homologous, though reduced, pterygoid dorsal process extends dorsally above the pterygoid condyle of the quadrate along the medial base of the orbital process into the orbital fossa in paleognaths like the ostrich and emu (*Figure 3—figure Supplement 1f*) (*McDowell, 1948; Zusi and Livezey, 2006*). The leaf-like palatine process is constricted along the middle third of the shaft and then expands mediolaterally rostrally. The caudal end of the palatine process extends caudally well beyond the level of the cotyle for the basipterygoid process, but the corresponding portion is reduced in IVPP V12707 and non-avialan theropods (*Figure 3—figure Supplement 1*) (*Ostrom, 1969; Barsbold and Osmólska 1999; Wang et al., 2021a*).

**Figure 3.**
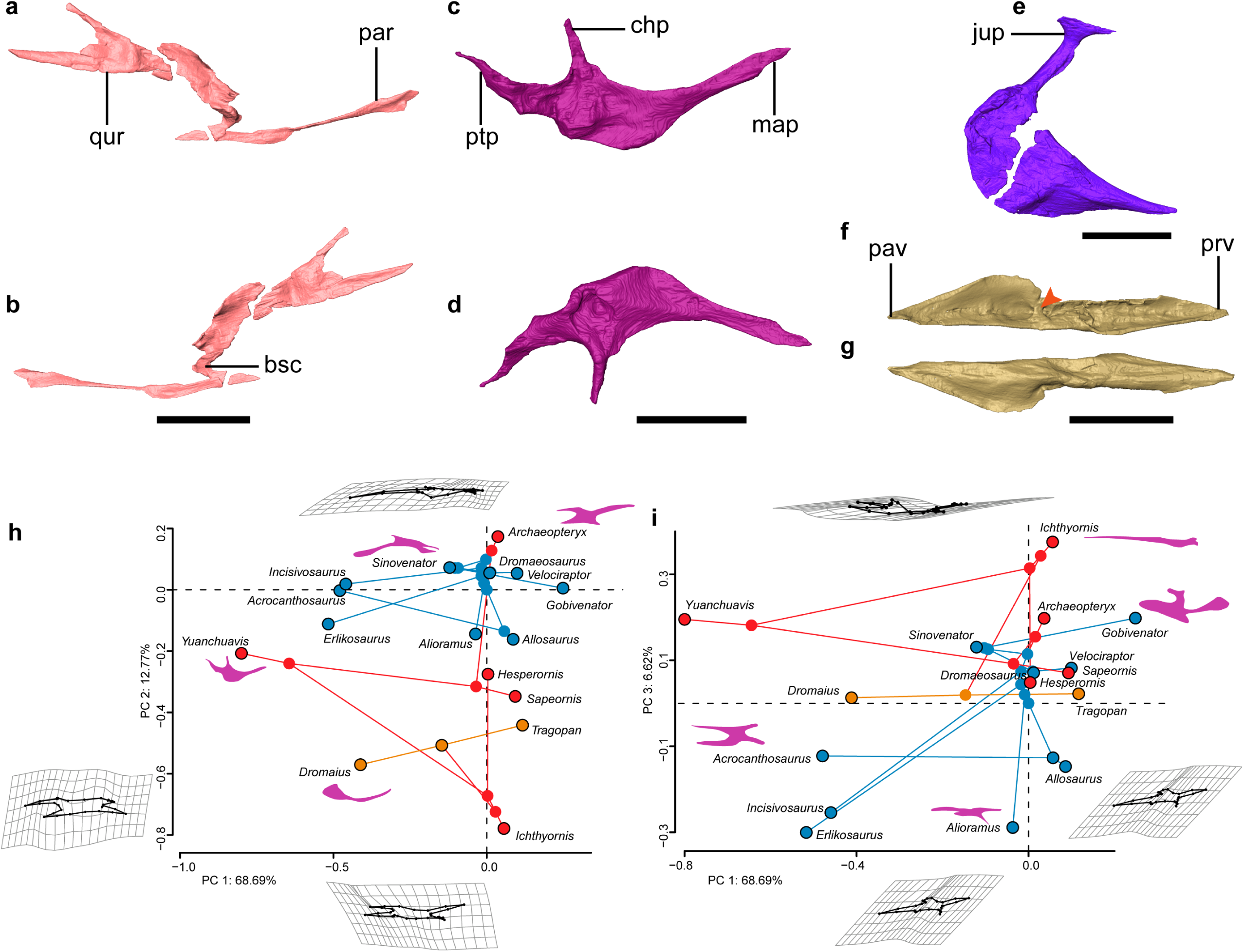
Palate anatomy of *Yuanchuavis*. **a–f**: Digital reconstruction of the right pterygoid in lateral (**a**) and ventral (**b**) view; right palatine in dorsal (**c**) and ventral (**d**) view; right ectopterygoid in dorsal view (**e**); and left vomer in dorsal (**f**) and ventral (**g**) view (arrowhead denotes the dorsal transverse ridge). **h, i**, Phylomorphospace showing the diversity of palatine shape in early diverging avialans and their close non-avialan theropod relatives based on the first three principal components (PC1–PC3), with deformation grids and wireframes from average to extreme; line drawings of the palatine in dorsal/ventral view are placed nearby corresponding taxa (blue circles: non-avialan theropods; red circles: Mesozoic avialans; orange circles: crown birds). bsc, basipterygoid process cotyle; chp, choanal process; jup, jugal process; map, maxillary process; par, palatine ramus; pav, palatine ramus of vomer; prv, premaxillary ramus of vomer; ptp, pterygoid process; qur, quadrate ramus. Scale bars, 10 mm (**a**–**d, f, g**), 2 mm (**e**). **Figure supplementary 1.** Comparison of pterygoid morphology. **Figure supplementary 2.** Comparison of palatine morphology. **Figure supplementary 3.** Comparison of vomer morphology. **Figure supplementary 4.** Diversity of palatine shape in early diverging avialans and non-avialan theropods. **Figure supplementary 5.** Landmark scheme.

While knowledge of the morphology of the palatine has been elusive among early avialans, the bone is well-preserved in *Yuanchuavis* (*Figure 3c, d*). As in *Falcatakely* (*O’Connor et al., 2020*), the bone is triradiate and lacks a jugal process present in *Archaeopteryx* and non-avialan dinosaurs (*Figure 3—figure Supplement 2a–f*) (*Ostrom, 1969; Barsbold and Osmólska, 1999; Mayr et al., 2007; Kundrát et al., 2019*). A jugal process is not preserved in the early pygostylian *Sapeornis* (*Figure 3—figure Supplement 2b*), which was interpreted as preservational bias (*Hu et al., 2019*). Given the shared condition and the exquisite preservation in *Yuanchuavis* and *Falcatakely*, we posit that the loss of the jugal process is genuine in these taxa, and represents a derived condition otherwise widely distributed among crownward ornithuromorphs (*Figure 3— figure supplement 2c, g, h*) (*Elzanowski, 1991; Elzanowski and Wellnhofer, 1996; Zusi and Livezey, 2006*). As in *Sapeornis* and *Hesperornis* (*Elzanowski, 1991; Hu et al., 2019*), the maxillary process is longer than the pterygoid process, in contrast to the condition in *Archaeopteryx* (*Figure 3—figure Supplement 2*) (*Elzanowski and Wellnhofer, 1996*). As in *Sapeornis* and *Falcatakely*, the caudally tapering pterygoid process is caudomedially directed with respect to the maxillary process, rather than being in line with the latter as in *Hesperornis* (*Elzanowski, 1991; Elzanowski and Wellnhofer, 1996*), and some non-avialan theropods (e.g., *Velociraptor*) (*Barsbold and Osmólska, 1999*). As in *Sapeornis* (*Hu et al., 2019*), the choanal process projects medially, rather than being hooked rostrally as in other early avialans (*Elzanowski, 1991; Elzanowski and Wellnhofer, 1996; O’Connor et al., 2020*), and some non-avialan theropods (*Currie, 1995; Barsbold and Osmólska, 1999*).

The ectopterygoid has a hooked jugal process that is expanded rostrocaudally at the lateral end, forming a large contact facet for the jugal, as in *Sapeornis* and some non-avialan theropods like *Allosaurus* (*Figure 3e*) (*Madsen, 1976; Hu et al., 2019*). In contrast, a similar expansion is absent among other early avialans such as *Archaeopteryx* and IVPP V12707 (*Elzanowski and Wellnhofe,r 1996; Wang et al., 2021a*). The main body of the ectopterygoid is fan-shaped, and its rostral end projects more rostrally than the state in *Archaeopteryx* and *Sapeornis* (*Mayr et al., 2005; Hu et al., 2019*).

The vomers are separated from each other as in the juvenile enantiornithine IVPP V127071 (*Figure 3f, g*; *Figure 3—figure Supplement 3a, b*), suggesting that the lack of fusion in the latter cannot be unequivocally attributed to ontogeny given that the holotype specimen here is from an osteologically mature individual. However, fused vomers are present in *Gobipteryx*, *Sapeornis*, and most crown birds (*Baumel and Witmer, 1993; Chiappe et al., 2001; Hu et al., 2019*) (*Figure 3—figure Supplement 2c*), and that variation demonstrates species-specific development of this element. The vomers differ from that of other known Mesozoic avialan and non-avialan theropods in that each vomer is concave dorsally with a thickened medial margin, and a dorsal transverse ridge is present at midshaft, dividing the vomer into premaxillary and pterygoid rami (*Figure 3f, Figure 3—figure Supplement 2*) (*Ostrom, 1969; Elzanowski and Wellnhofer, 1996; Lautenschlager et al., 2014; Hu et al., 2019; Wang et al., 2021a*). The premaxillary ramus is spear-shaped in dorsal view. The pterygoid ramus reaches its maximum mediolateral width at its midpoint and narrows caudally. Unlike the mediolaterally compressed form observed in IVPP V12707 (*Wang et al., 2021a*), the pterygoid ramus is dorsoventrally compressed and forms a dorsal groove to articulate with the pterygoid. A caudodorsal process, projecting from the caudal end of the vomer like that of *Sapeornis* and some non-avialan theropods (*Hu et al., 2019*), is absent.

The occipital elements around the foramen magnum are compressed, but the occipital condyle is distinct with a heart shaped outline and a nerve foramen on the left side of its base. The blunt though pointed basipterygoid process is pronounced as in some enantiornithines (e.g., *Zhouornis*, *Brevirostruavis*) (*Zhang et al., 2013; Li et al., 2021*), paleognaths (*Gussekloo and Bout, 2005*), and non-avialan theropods (*Holliday and Witmer 2008*) (*Figure 1, Figure 1—figure Supplement 1*).

As in other enantiornithines (*O’Connor and Chiappe, 2011*), the dentary has parallel dorsal and ventral margins and a caudoventrally sloping caudal margin (*Figure 2m, n*). Seventeen dentary teeth were present on each side (11 teeth in situ and six alveoli; *Figure 2—figure Supplement 1f)*. Meckel’s groove extends rostrally to the level of the third dentary tooth and appears to be fully covered medially by the triangular splenial (*Figure 2—figure Supplement 1f, g*). The post-dentary mandibular elements are compressed with each other (or fused partially), preventing digital segmentation of the individual elements (*Figure 2o*). Unlike IVPP V12707 (*Wang et al., 2021a*), neither a mandibular fenestra nor a coronoid process are present. There appears to have been a small retroarticular process in *Yuanchuavis*, but a prominent one is present in some enantiornithines such as bohaiornithids and *Brevirostruavis* (*Wang et al., 2014; Stidham and O’Connor, 2021*). The caudal end of the right jaw is broken just caudal and ventral to its articulation with the quadrate, but the preserved base of the retroarticular process does extend somewhat dorsally. The caudal end of the left jaw appears possibly complete and exhibits an overall squared caudal end suggesting the complete shape of the process.

The atlas has a complete fused neural arch, rendering this element ring-like in cranial view (*Figure 1—figure Supplement 2*). The condyloid fossa is subcircular in outline and bears a shallow incisure fossa. A pair of short costal processes project caudodorsally as in some enantiornithines such as *Piscivorenantiornis* (*Wang and Zhou, 2017*). The articular facet for the axis is mediolateral wider than dorsoventrally high. The elongate axis is craniocaudally longer than dorsoventrally high. The caudal articular facet of the centrum seems to be heterocoelic, being mediolaterally convex and dorsoventrally concave, as in crown birds.

## Discussion

The exquisitely preserved skull of *Yuanchuavis*, visualized through high-resolution x-ray computed tomography, enables reconstruction of the cranial morphology of early avialans with great fidelity. In particular, this reconstruction includes elements that are otherwise rarely preserved or poorly known, but are key to the analysis of macroevolutionary and functional properties of the skull (*Zusi, 1993; Sander et al., 2001*). Given the presence of pointed and lightweight jaw bones, enlarged orbit, and vaulted cranial roof, the skull is clearly bird-like. However, the morphologies of the temporal and palatal regions speak to its archetypal nature, and attest to the retention of typical non-avialan dinosaurian conditions with little modifications. The combination of a T-shaped postorbital with an elongate jugal process and a large squamosal with a prominent postorbital process indicate the presence of postorbital and temporal bars that completely separate the supratemporal and infratemporal fenestrae, and the orbit from each other (*Figure 2p*). In addition, the infratemporal fenestra likely is separated from the quadrate fenestra given the length of the squamosal process of the quadratojugal and the quadratojugal process of the squamosal. These observations unambiguously show that *Yuanchuavis* has a diapsid skull, which so far has been confirmed in *Archaeopteryx* (*Elzanowski and Wellnhofer, 1996; Mayr et al., 2005*), the early pygostylian *Sapeornis* and confuciusornithiforms (*Chiappe et al., 1999; Hou et al., 1999*), and two enantiornithines (*Longusunguis* and IVPP V12707) among Cretaceous avialans (*Wang et al., 2021a*), showing that this ancestral temporal configuration is conserved evolutionarily well into the early avialan diversification; whereas the recently reported Late Cretaceous enantiornithine *Yuornis* shows the independent loss of the postorbital bar along with reduction of other cranial elements (*Xu et al., 2021*).

The structure of the palate is functionally vital to the feeding behaviors of vertebrates (*Zusi and Livezey, 2006; Holliday and Witmer, 2008*), and it underpins the diverse forms of cranial kinesis present in crown birds (*Zusi, 1984; Gussekloo et al., 2017*). However, our understanding of the transformation of the palate from the ancestral heavily built and immobile dinosaurian morphology into the flexible derived crown bird condition has been hindered by a lack of detailed early avialan skull records (*Witmer and Martin, 1987; Field et al., 2018a; Hu et al., 2019*). Reconstruction of the complete palate of *Yuanchuavis* reveals a stout vomer that is dorsoventrally compressed and likely forms a large overlapping contact with the maxilla/premaxilla as in *Sapeornis* and non-avialan theropods (*Figure 3, Figure 3—figure Supplement 3*) (*Ostrom, 1969; Holliday and Witmer, 2008*). The pterygoid is morphologically nearly identical to that of non-avialan theropods in having a dorsally oriented quadrate ramus that extends medially bypassing the orbital process of the quadrate, resulting in a scarf joint widely distributed among non-avialan dinosaurs (*Holliday and Witmer, 2008; Tsuihiji et al., 2014*). The forked condition of this quadrate ramus is shared with close relatives of Avialae like *Linheraptor*, and this dinosaurian pterygoid previously has been recognized only among early avialans in the juvenile enantiornithine IVPP V12707 (*Wang et al., 2021a*). The presence of this plesiomorphic morphology in both a juvenile (i.e., IVPP V12707) and a skeletally mature (here, *Yuanchuavis*) enantiornithine eliminates ontogenetic variation as a potential source for its morphology, and signifies conclusive evidence as to the presence of the kind of dinosaurian pterygoid in enantiornithines, along with its functional implications.

The palatine exhibits a peculiar morphology distinguishable from that of other early diverging avialans and non-avialan theropods in having a medially directed choanal process, having a caudomedially oriented pterygoid process, and lacking a jugal process. To explore evolutionary changes in palatine morphology across the theropod to bird transition, we performed landmark-based geometric morphometric analyses. A phylomorphospace was created by using the first three principal components (PCs 1**–** 3: >85% of shape variances) calculated from generalized Procrustes superimposition and followed by phylogenetic principal components analysis to account for phylogenetic non-independence (*Figure 3h, i; Figure 3—figure Supplement 4*). PC1 is aligned most with the length of the choanal process, and the length of the medial distance defined by the choanal-pterygoid processes relative to the lateral distance defined by the jugal-maxillary processes, with *Yuanchuavis* exhibiting the lowest score (*Figure 3h*). PC2 corresponds to the length of the jugal process and the overall shape of the palatine, and differences along this axis distinguishes avialans from *Archaeopteryx* and non-avialan theropods, reflecting the absence of the jugal process in avialans crown-ward of *Archaeopteryx*. PC3 mainly describes the slenderness of the palatine, and early diverging theropods (e.g., *Allosaurs*, *Erlikosaurus*) with rectangular palatines clustered together (*Figure 3i*). *Yuanchuavis* is widely separated from other stem and crown avialans, and representatives of major non-avialan theropods in palatine morphospace (*Figure 3h, i; Figure 3—figure Supplement 4*). This preliminary geometric morphometric analysis suggests that the loss of the jugal process represents the major structural change of the palatine in early bird evolution. The release of constraints imposed by contacts with other elements, combined with selection for derived feeding behaviors and functions, likely guided the morphological and functional divergence of the palatine. Future studies with increased species sampling are needed to further test this hypothesis.

With >10,000 species, neognaths are the most morphologically and ecologically diverse clade of modern amniotes (*Gill, 2007*), with much of their success attributed to key evolutionary novelties such as powered flight, but also their uniquely kinetic crania (*Zusi, 1993; Gussekloo and Bout, 2005; Lovette and Fitzpatrick, 2016*). Avian cranial kinesis has been demonstrated to have improved feeding performance by increasing biting force, jaw closing speed, and food handling precision (*Bock, 1964; Zusi, 1984*). Avian cranial kinesis works via pathways comprised of two kinetically permissive linkage systems: the quadrate-quadratojugal-jugal-rostrum on the lateral margin, and the quadrate-pterygoid-palatine-vomer on the palatal aspect, which coordinate in transmitting force and movement of the musculature to the rostrum through the sliding movement of the entire palate (*Figure 4*) (*Bock, 1964; Holliday and Witmer, 2008; Bhullar et al., 2016; Wang et al., 2021a*). However, to achieve this movement, the involved elements have been radically transformed from the primitive condition in non-avialan theropods, in which these cranial bones are robust and articulate with each other through immobile and largely sutural contacts, to lighter elements with condylar or otherwise reduced contacts (*Figure 4a, b*) (*Holliday and Witmer, 2008*). Our study demonstrates that neither of these two pathways utilized in modern cranial kinesis are present or functional in *Yuanchuavis*. The rostrocaudal movement of the jugal bar to protract/retract the upper jaw is prevented by the complete temporal bars and the bracing of the ectopterygoid. The sliding movement of the palatal elements is severely restricted by the prominent basipterygoid process (that fits into a cotyle on the pterygoid) and the lack of permissive synovial articulations between the quadrate-pterygoid and pterygoid-palatine contacts (*Figure 4c, d*).

**Figure 4.**
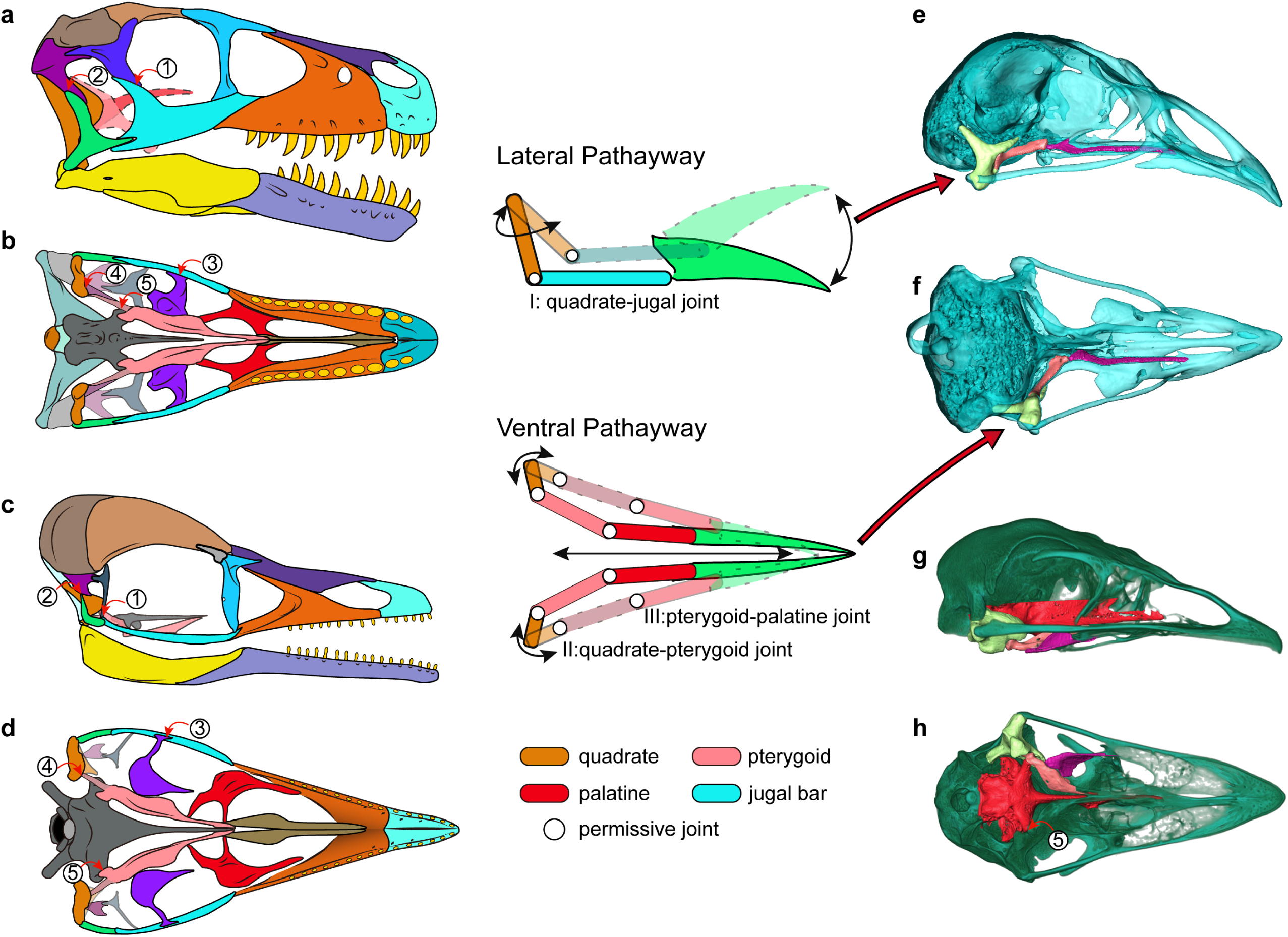
Cranial kinesis in early bird evolution. The typical avian cranial kinesis is realized through two pathways (schematic drawing in the middle): the quadrate-jugal bar-rostrum on the lateral side, and quadrate-pterygoid-palatine on the ventral side, highlighted in the galliform *Tragopan caboti* (**e, f**). These two pathways are restricted by the postorbital bar (1), squamosal-quadratojugal contact (2), ectopterygoid (3), the scarf join between quadrate-pterygoid (4), and the prominent pointed basipterygoid process (5) in non-avialan theropods (**a, b**) and enantiornithines (**c, d**), indicating an akinetic skull. The ventral pathway is also absent in most paleognaths (**h**: at least restricted by non-permissive contact 5). **a–d**, Skull reconstructions of *Dromaeosaurus* (Theropod: Dromaeosauridae) in lateral (**a**) and ventral (**b**) view (modified from *Currie, 1995*), *Yuanchuavis* (Avialae: Enantiornithes) in lateral (**c**) and ventral (**d**) view. **e–g**, Digital renderings of skulls of *Tragopan caboti* (Galloanserae: Galliformes) in lateral (**e**) and ventral (**f**) view, and *Dromaius novaehollandiae* (Palaeognathae: Casuariiformes) in lateral (**g**) and ventral (**h**) view.

However, as noted previously (*Wang et al., 2021a*), the presence of the scarf joint between the quadrate and pterygoid along with the potential rotation of the quadrate mediolaterally around its long dorsoventral axis as in crown birds (*Zusi, 1984*) may have allowed for some rudimentary force transmission from the quadrate rostrally along the palatal bone contacts with the basipterygoid process acting as a fulcrum. That hypothesized movement among early avialans may have been the functional foundation from which other features were evolved and exapted to complete the mosaic of modern avian cranial kinesis. While it would be easy to think of the modification of the ancestral dinosaur or theropod skull as having been driven towards kinesis (i.e., the idea of preadaptation (*Gould and Vrba, 1982*)), many of these features evolved in akinetic avialan skulls and clearly were selected for a function other than modern avian cranial kinesis. These features were only later exapted for kinetic function. Our study confirms that the akinetic skull present in earliest diverging avialans was retained in enantiornithines (*Wang et al., 2021a*). Despite their akinetic origins, features such as the loss of the jugal process of the palatine and further reduction in the bones of the temporal region would go on to be exapted and modified further as part of the evolutionary assembly of kinetic skulls. Many of the skeletal modifications necessary for cranial kinesis were pieced together within the crownward clade Ornithuromorpha (*Witmer and Martin, 1987; Wang et al., 2021a*), with the oldest known records of the neognath style of palatine-pterygoid contact, the condylar contact between the quadrate and pterygoid, and loss of the diapsid temporal condition in the Late Cretaceous *Ichthyornis* and *Hesperornis* (*Elzanowski, 1991; Field et al., 2018a*)

The mixture of plesiomorphic temporal and palatal regions and derived facial anatomies captured in this single enantiornithine skull vividly demonstrates how the avialan cranium has been shaped deeply by evolutionary mosaicism—a hypothesis proposed by studies involving crown birds but still awaiting paleontological evidence (*Bhullar et al., 2016; Felice et al., 2020; O’Connor et al., 2020; Plateau and Foth, 2020*). That difference is even more conspicuous between the skull and the postcranial skeleton of *Yuanchuavis*, given the presence of derived avialan features such as heterocoelic cervical vertebrae (Figure 1—*figure supplement 2*), a fused synsacrum and pygostyle, and a reversed hallux (*Wang et al., 2021b*), demonstrating that the avialan bauplan is highly modular (*Orkney et al., 2021*). This evolutionary modularity may help to explain the counterintuitive fact that while enantiornithines apparently were constrained by the retention of ancestral akinetic dinosaurian cranial configurations, they were the dominant avialan clade over a large part of the Cretaceous around the global with their enhanced locomotion (powered flight) and diverse feeding strategies conveyed by disparate rostrum morphologies, hyoid, and dental morphologies (*O’Connor and Chiappe, 2011; O’Connor et al., 2020, 2011; Li et al., 2021*). Comparing the extinction of enantiornithines at the end of the Cretaceous to the survival of members of their sister clade Ornithuromorpha (*Field et al., 2018b*), it is easy to hypothesize that the ancestral akinetic avialan skull, despite its successful use among avialans throughout the Cretaceous, did not allow for the adaptations or extreme selection afforded by the fully kinetic skulls of crown birds. That kinesis with its origin spread across various points of the Mesozoic avialan tree has contributed to the greatest radiation of avialans and their occupation of diverse habitats around the world.

## Methods

### Revised systematic paleontology of *Yuanchuavis kompsosoura*

Avialae Gauthier 1986

Ornithothoraces Chiappe 1995

Enantiornithes Walker 1981.

Pengornithidae Wang et al. 2014.

Yuanchuavis kompsosoura Wang et al. 2021.

Revised diagnosis. A large pengornithid enantiornithine that is distinguishable from other pengornithids on basis of following features (*autapomorphy): premaxilla bearing six teeth with its rostral tip edentulous*; lacrimal bearing a lateral flange; dorsoventrally broad orbital process of the quadrate; unfused vomers bearing a transverse dorsal ridge*; palatine having a medially directed choanal process and lacking a jugal process; pterygoid with a forked quadrate ramus; cranial thoracic vertebrae having well-developed hypapophyses; middle thoracic centra laterally excavated by broad fossae; and pygostyle with elongate ventrolateral processes and short dorsal processes (modified from *Wang et al., 2021b*).

### X-ray computed tomography imaging

To optimize the scanning resolution, the skull and the two cranial vertebrae (atlas and axis) of the holotype of *Yuanchuavis kompsosoura* (IVPP V27883) were isolated from the slab. The skull was scanned using the industrial CT scanner Phoenix v-tome-x at the Key Laboratory of Vertebrate Evolution and Human Origins, Institute of Vertebrate Paleontology and Paleoanthropology (IVPP, Beijing, China), with a beam energy of 140 kV and a flux of 150 μA at a resolution of 11.68 μm per pixel. The resulting scanned images were imported into Avizo (version 9.2.0) for digital segmentation, rendering, and reconstruction. The obtained three-dimensional models were optimized in MeshLab (version 2012.12). CT data of the enantiornithine IVPP V12707, and two modern birds (*Tragopan caboti* and *Dromaius novaehollandiae*) produced during our previous research were re-analyzed and rendered for comparison (*Wang et al., 2021a*).

### Geometric morphometric analysis

In order to trace changes of palatine morphology among early diverging avialans and their close non-avialan theropod relatives, a landmark-based geometric morphometric analysis was performed. The palatine is generally poorly preserved in early avialan and non-avialan theropods, and we only have been able to assemble a dataset of 14 fossil taxa that preserve this element intact, including nine non-avialan theropods and five Mesozoic avialans (*Supplementary file 1*). Two crown birds—*Dromaius novaehollandiae* (Palaeognathae: Casuariiformes) and *Tragopan caboti* (Neognathae: Galliformes)—were included for comparison. Despite the small sample size, the dataset contains species from the major clades of non-avialan theropods and early avialans, and therefore should encompass the morphological disparity of palatine with moderate sufficiency. Seventeen landmarks and one curve (15 evenly spaced semi-landmarks along the lateral margin of the palatine) were digitized on the two-dimensional geometry of the palatine in dorsal/ventral aspect using the tpsDIG software (see *Figure 3—figure supplement* 5 and *Supplementary file 2* for landmark schemes) (*Rohlf, 2009*). A three-dimensional geometric morphometric analysis was not employed here, because this method would reduce the sample size greatly and add little additional information considering that the palatine is a dorsoventral sheet-like element in most taxa. To remove the non-biological effects of rotation, scaling and translation (*Zelditch et al., 2012*), the raw coordinate data was transformed by generalized Procrustes analysis (GPA), with the semi-landmarks slid to minimal bending energy, using the gpagen function in the R package geomorph (version 4.0.0) (*Adams and Otárola-Castillo, 2013*). To account for the non-independence of phenotypes among taxa because of shared history, a phylogenetic principal components analysis (pPCA) was performed (*Revell, 2009*). First, a super tree containing only the focal taxa was compiled with reference to recent phylogenetic work (*Turner et al., 2012; Wang et al., 2021b*). This phylogeny was time-calibrated using tip dates bracketed by the first and last appearance datum of the geological stages or epochs in which each taxon was collected (*Brusatte, 2011*). Zero-length branches were smoothed using the “minimum branch length” embedded in the timePaleoPhy function in the R package paleotree (*Bapst, 2012*). Then, the GPA transformed coordinate data and time-calibrated phylogeny were subjected to pPCA using the gm.prcomp function in the R package geomorph, which performed a generalized least square estimation of a covariance matrix (*Collyer and Adams, 2021*). A phylomorphospace was constructed using the principal components scores of taxa and ancestral nodes to visualize changes of palatine geometry along line to early avialans (*Figure 3h, i*).

In addition, a traditional geometric morphometric analysis also was applied for comparison (*Figure 3—figure Supplement 4*). The first three PCs account for 83.51% of the shape variance. PC1 corresponds to the relative length of the jugal process, and differences along this axis distinguish all avialans except *Archaeopteryx* from non-avialan theropods (reflecting the presence/absence of the jugal process). PC2 is most aligned with the discrepancy between the medial distance defined by the choanal and pterygoid processes and the lateral distance defined by the maxillary and jugal processes, with *Yuanchuavis* exhibiting the highest scores. PC3 describes the degree of mediolateral curvature of the palatine body, and derived ornithuromorphs and *Yuanchuavis* are positioned on the positive side along this axis.

## Acknowledgements

We thank Song Miao and Jiutong Feng for helping in CT data scanning and segmentation. We thank Han Hu for sharing photographs of *Sapeornis*. This research is supported by the Key Research Program of Frontier Sciences, CAS (ZDBS-LY-DQC002), the National Natural Science Foundation of China (42288201, 42172029), and the Tencent Foundation (through the XPLORER PRIZE).

## Author Contributions

Min Wang conceived the study; Min Wang conducted the digital reconstruction of the specimen; and Min Wang, Thomas A. Stidham, Jingmai K. O’Connor, and Zhonghe Zhou wrote the manuscript.

## Data availability

The specimen (IVPP V27883) described in this study is archived and available on request from the Institute of Vertebrate Paleontology and Paleoanthropology (IVPP), Chinese Academy of Sciences, Beijing, China. Additional figures of cranial anatomy are available in the Supplementary Information. The three-dimensional models (STL) are archived and available on Dryad (https://datadryad.org/stash/share/dYr1L9bLrtkO8WuevUVcDj7gnJ0W-9gv7ZiD1iv7CUU), or from the corresponding authors.

## Legends for supplementary figures

**Figure 1—figure supplement 1.**
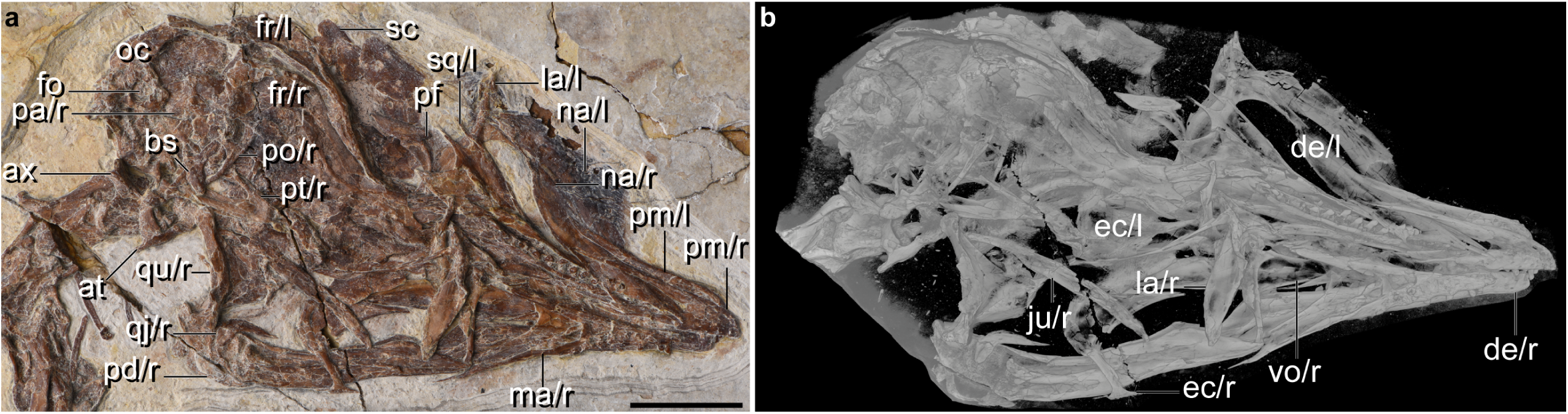
Cranial anatomy of *Yuanchuavis*, IVPP27883. **a,** Photograph. **b,** CT imaging. ax, axis; at, atlas; bs, basipterygoid process; de, dentary; ec, ectopterygoid; fo, foramen magnum; fr, frontal; ju, jugal; la, lacrimal; ma, maxilla; na, nasal; oc, occipital region; pa, parietal; pf, prefrontal; pd, post-dentary mandible; pm, premaxilla; po, postorbital; pt, pterygoid; qj, quadratojugal; qu, quadrate; sc, sclerotic bone; sp, splenial; sq, squamosal; vo, vomer; r/l, right/left side. Scale bar, 10 mm.

**Figure 1—figure supplement 2.**
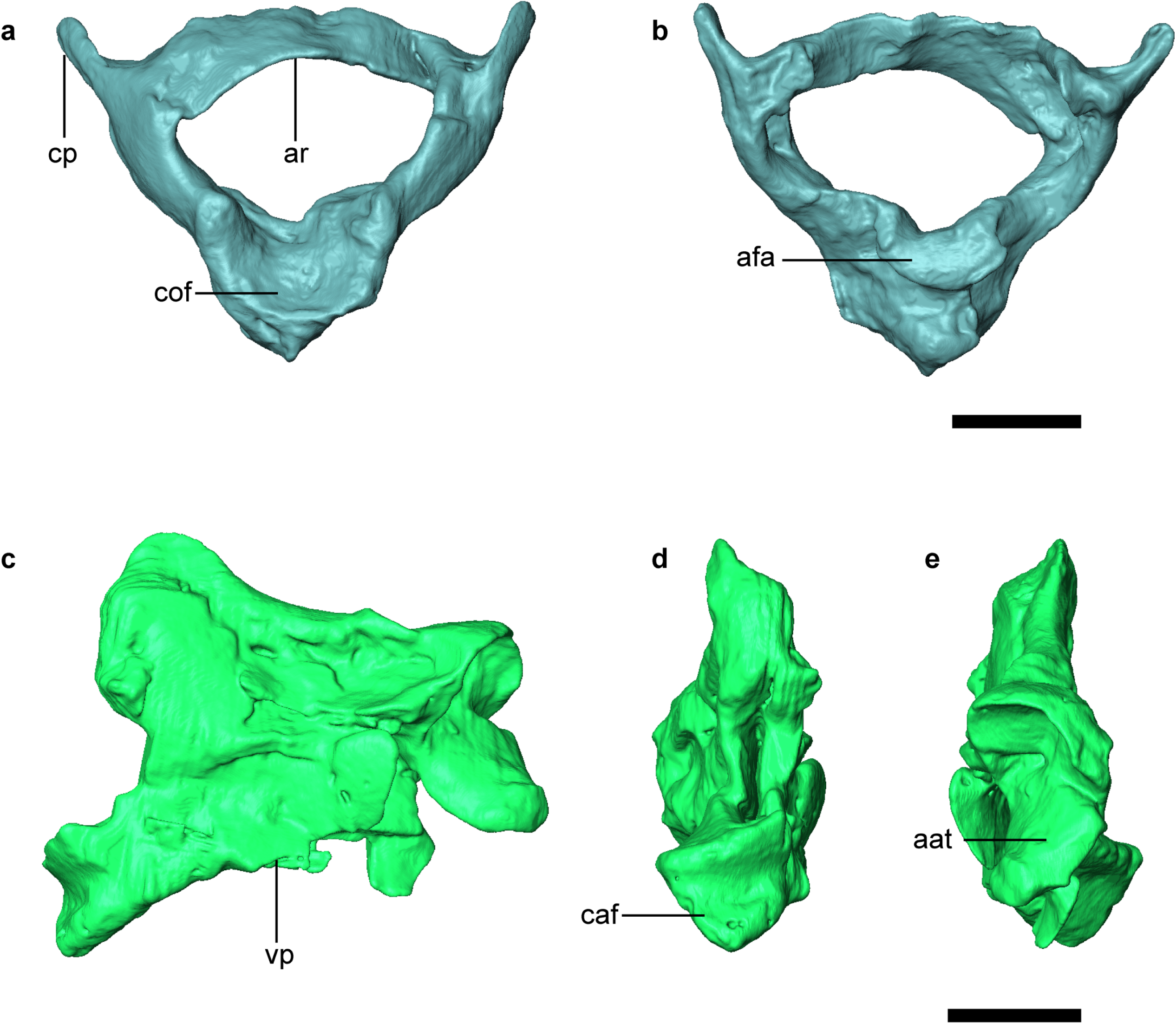
Digital reconstruction of atlas and axis of *Yuanchuavis*. **a, b,** Atlas in cranial (**a**) and caudal view (**b**). **c–e,** Axis in lateral (**c**), caudal (**d**), and cranial view (**e**). aat, articular facet for atlas; afa, articular facet for axis; ar, arcus atlantis; caf, caudal articular facet; cof, condyloid fossa; cp, costal process; tv, transverse process; Scale bars, 2 mm.

**Figure 2—figure supplement 1.**
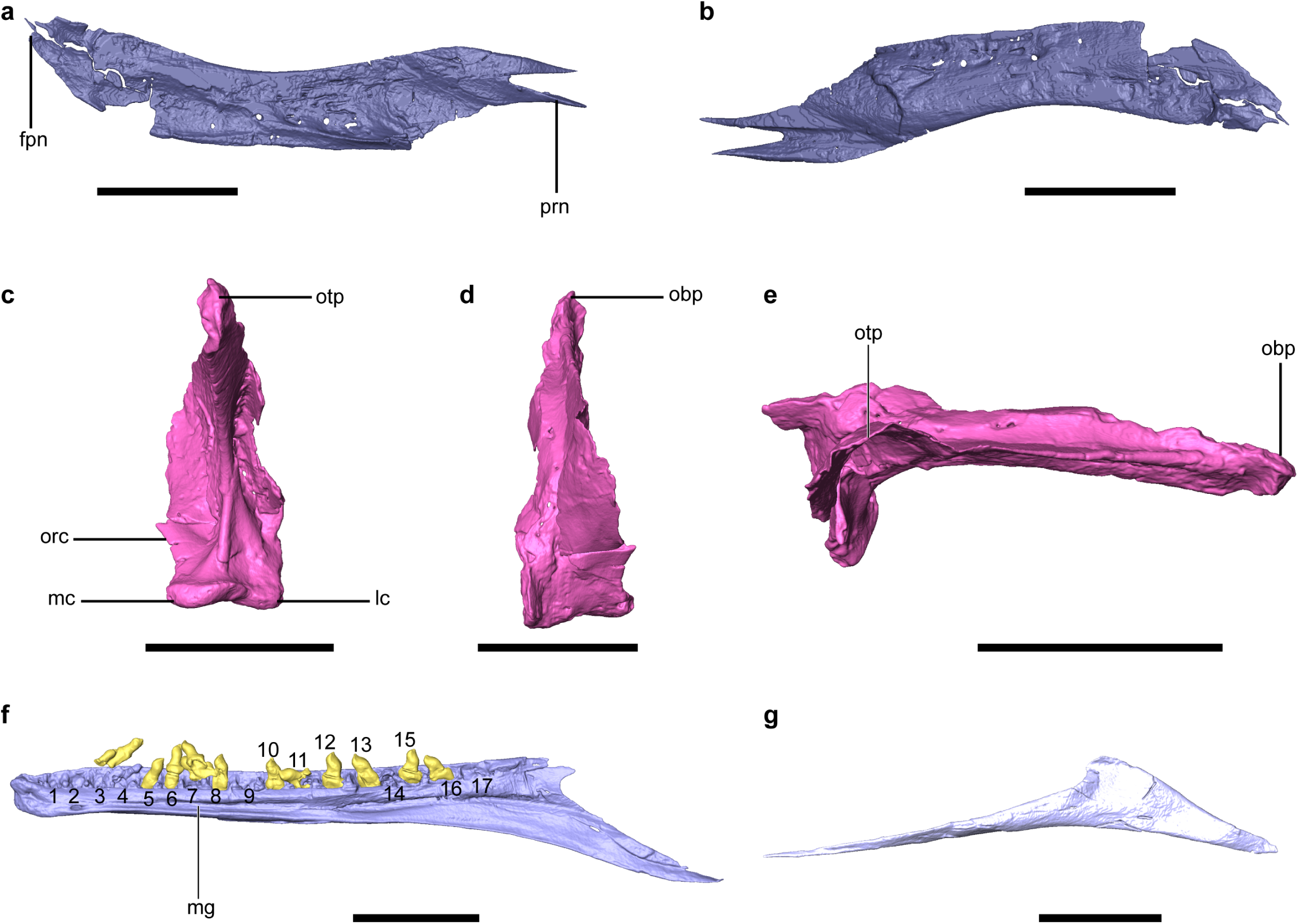
Additional cranial anatomy of *Yuanchuavis*. **a–g**, Digital reconstruction of the left nasal in medial (**a**) and lateral (**b**) view, right quadrate in caudal (**c**), rostral (**d**), and dorsal (**e**) view, right dentary in medial view (**f**), and right splenial in medial view (**g**). fpn, frontal process of nasal; lc, lateral condyle; mc, medial condyle; mg, Meckel’s groove; obp, orbital process; orc, orbitocondylar crest; otp, otic process; prn, premaxillary process of nasal; 1–17, dentary tooth count. Scale bars, 10 mm (**a, b, f, g**), 5 mm (**c–e**).

**Figure 3—figure supplement 1.**
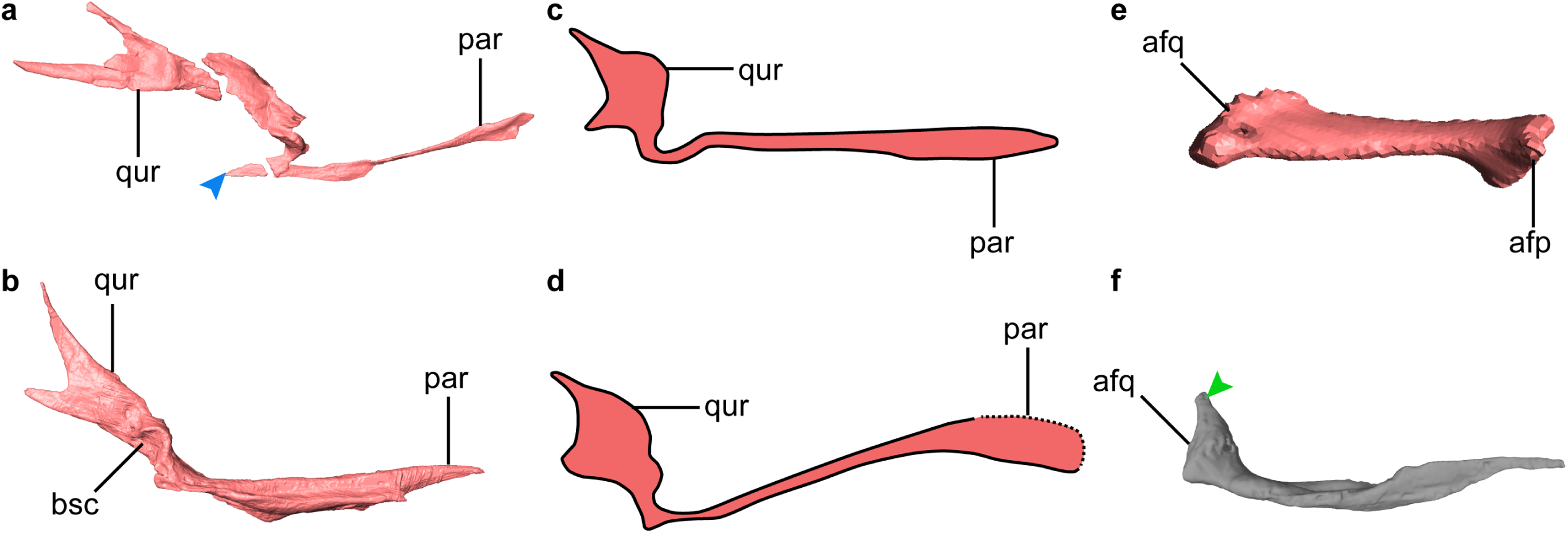
Comparison of pterygoid morphology. **a, b**, Digital reconstruction of the pterygoid of *Yuanchuavis* (**a**, right element in lateral view), and enantiornithine IVPP V12707 (**b**, left element in medial view). **c, d,** Line drawing of the pterygoid of *Sinornithosaurus millenii* (Theropod: Dromaeosauridae; left in medial view; **c**), and *Sinovenator changii* (Theropod: Troodontidae; left in medial view; **d**). **e, f,** Digital reconstruction of the pterygoid of *Tragopan caboti* (**e**, left element in ventrolateral view), and *Dromaius novaehollandiae* (**f**, left element in lateral view). afq, articular facet for quadrate; afp, articular facet for parasphenoid rostrum; bsc, basipterygoid cotyla; par, palatine ramus; qur, quadrate ramus. The blue arrowhead in (**a**) denotes the caudal extension that is absent in other enantiornithines and non-avialan theropods (**b–d**). The green arrowhead in (**f**) indicates the potential homologous, but reduced quadrate ramus present in some paleognaths. All figures are not scaled.

**Figure 3—figure supplement 2.**
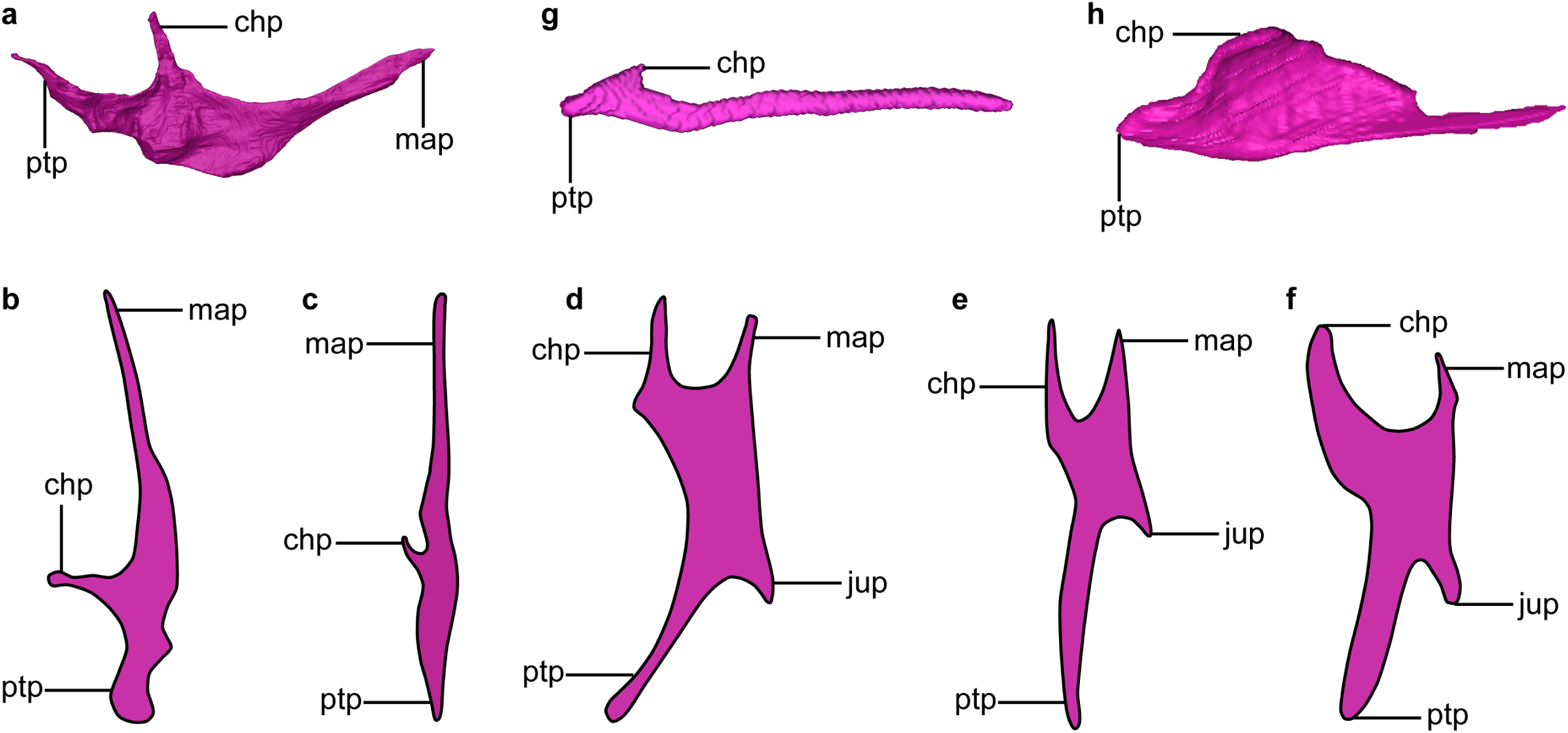
Comparison of palatine morphology. **a, g, h,** Digital reconstruction of the palatine of *Yuanchuavis* (**a**, right element in dorsal view), *Tragopan caboti* (**g,** left element in ventral view), and *Dromaius novaehollandiae* (**h**, left element in ventral view). **b–f,** interpretative line drawing of the palatine of selected early diverging avialans and non-avialan theropods (converted to right element in ventral view): *Sapeornis* (**b**), *Hesperornis* (**c**), *Archaeopteryx* (**d;** modified from *Bhullar et al., 2016*), *Velociraptor* (**e;** modified from *Barsbold and Osmólska, 1999*), and *Dromaeosaurus* (**f;** modified from *Currie, 1995*). chp, choanal process; jup, jugal process; map, maxillary process; ptp, pterygoid process. All figures are not scaled.

**Figure 3—figure supplement 3.**
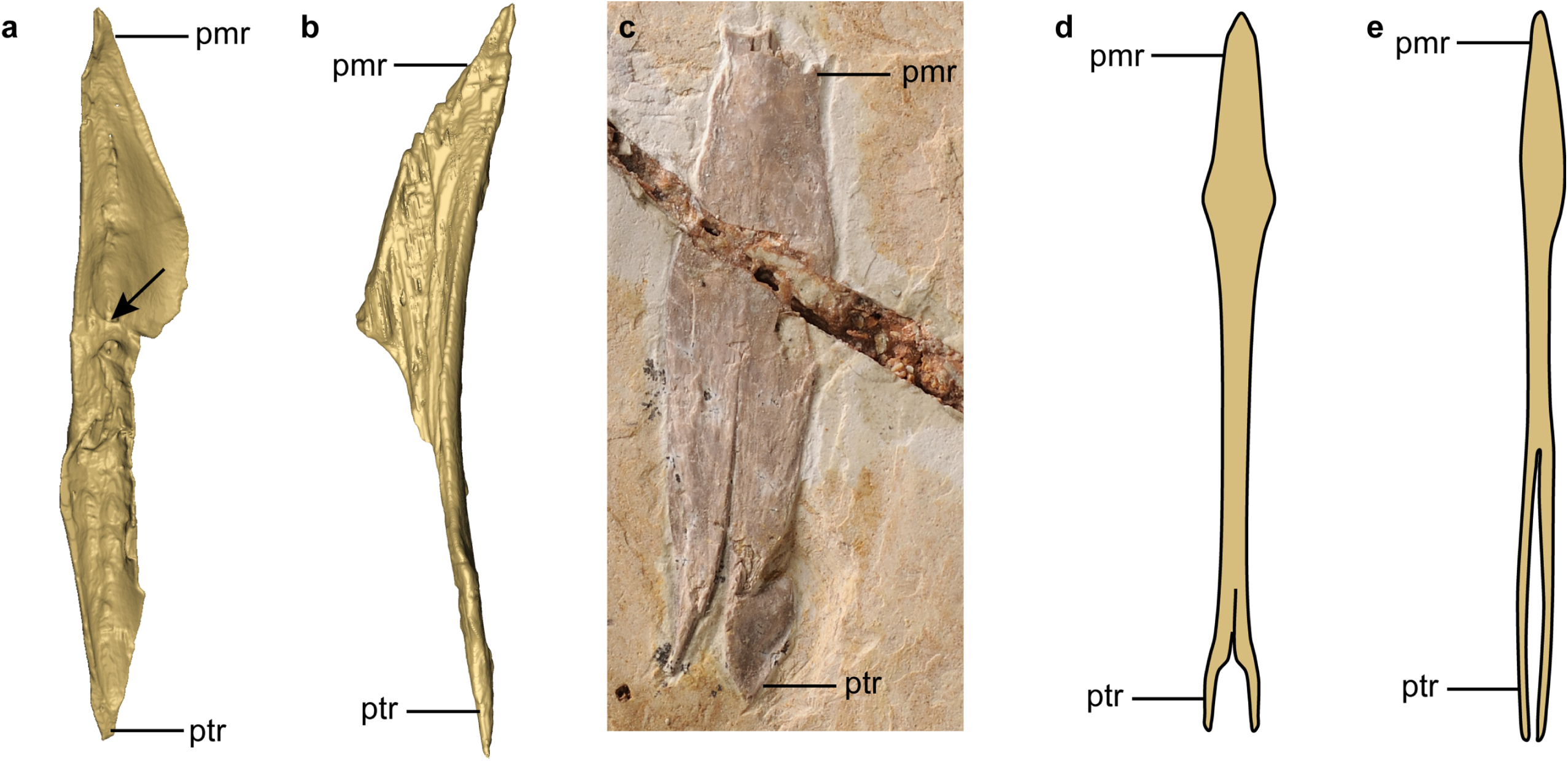
Comparison of vomer morphology. **a, b,** Digital reconstruction of the right vomer of *Yuanchuavis* (**a**), and left vomer of enantiornithine IVPP V127071 (**b**) in dorsal aspect. **c,** Photograph of *Sapeornis* (IVPP V19058). **d, e,** Line drawing the vomer of *Dromaeosaurus* (**d**) and *Allosaurus* (**e**) in dorsal view (modified from *Currie, 1995; Barsbold and Osmólska, 1999*). The rostral ends of the vomers are fused in (**c–e**), which are not in the two enantiornithines (**a, b**). pmr, premaxillary ramus; ptr, pterygoid ramus. The arrowhead in (**a**) denotes the transverse ridge that is absent in other taxa. All figures are not scaled.

**Figure 3—figure supplement 4.**
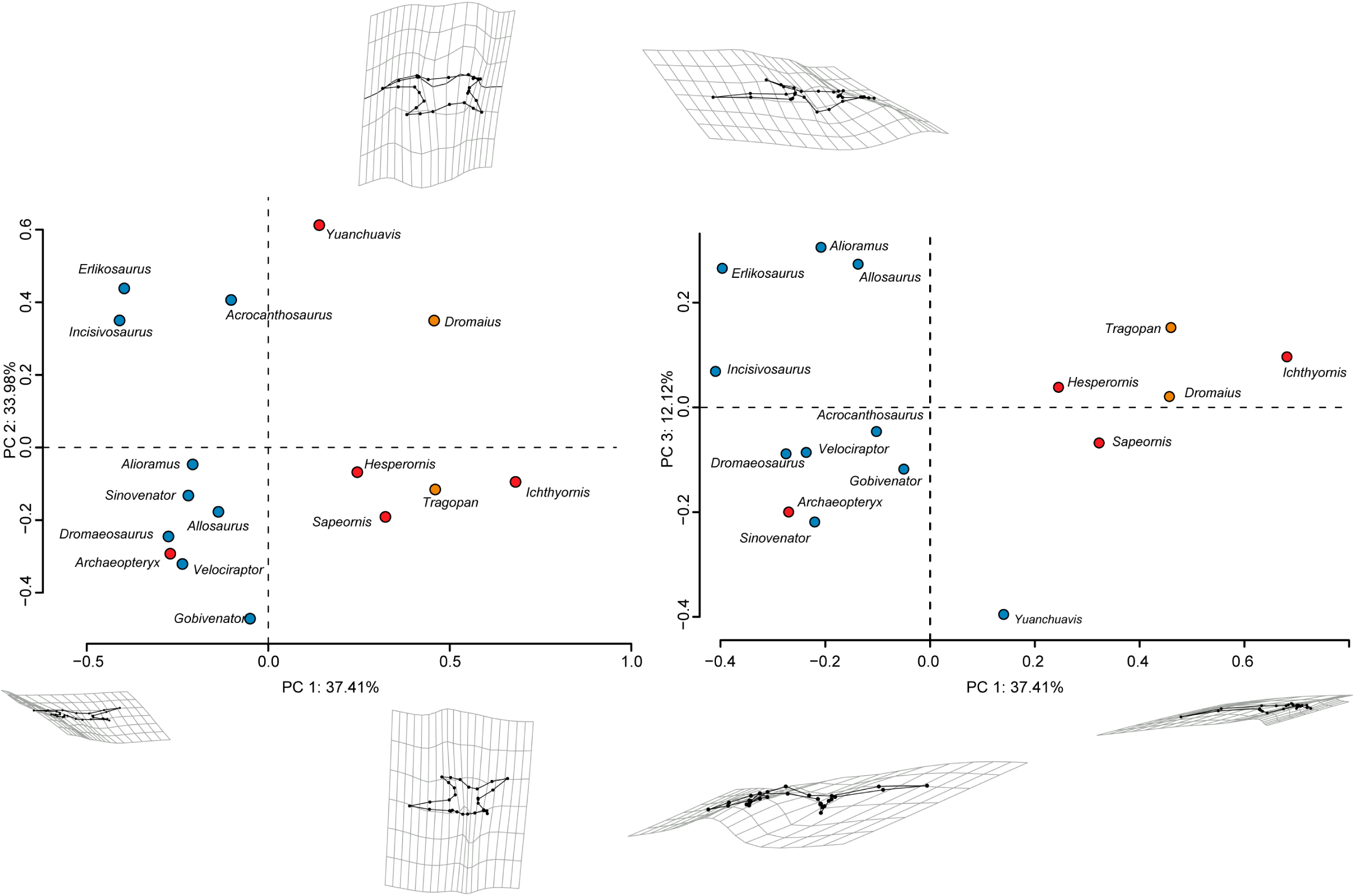
Diversity of palatine shape in early diverging avialans and non-avialan theropods. Morphospace based on the first three principal components (PC1–PC3), with deformation grids and wireframes from average to extreme (blue circles: non-avialan theropods; red circles: Mesozoic avialans; orange circles: crown birds).

**Figure 3—figure supplement 5.**
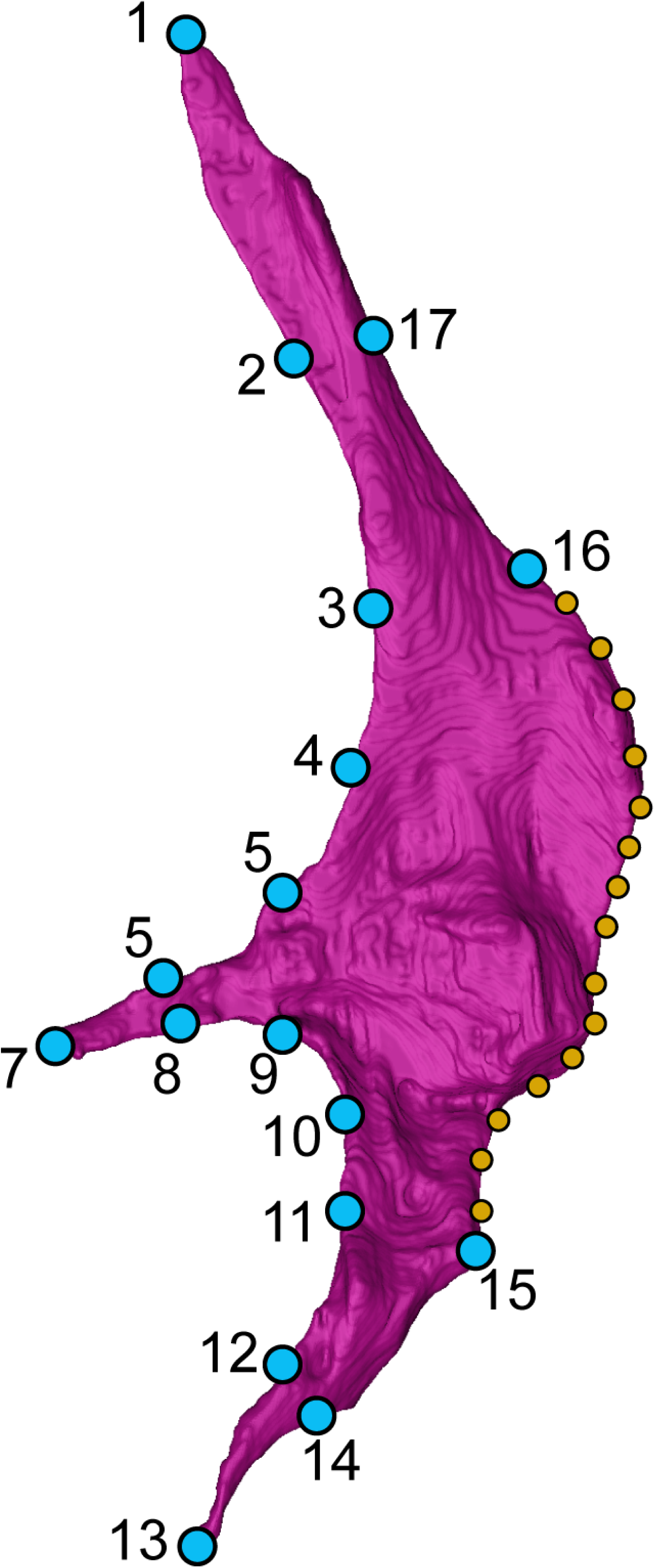
Landmark scheme. 17 landmarks (blue dots) and 15 semi-landmarks (orange dots) were digitized on the CT reconstructed right palatine of *Yuanchuavis* in dorsal view. See *Supplementary file 3* for anatomical description of landmarks and semi-landmark placements.

**Supplementary video 1. Three-dimensional digital model of *Yuanchuavis kompsosoura* with rotation around vertical axis.**

**Supplementary video2. Three-dimensional digital model of *Yuanchuavis kompsosoura* with rotation around horizontal axis.**

## Supplementary file 1

### Anatomical description of landmarks and semi-landmarks positions

Landmarks 1-17 (Figure 3—*figure supplement 5*):

Rostral tip of the maxillary process
Medial midpoint of the maxillary process
Medial deflection point between the maxillary process and palatine body
Midpoint of medial margin connecting the maxillary and choanal processes
Rostral deflection point between the choanal process and palatine body
Rostral midpoint of the choanal process
Rostral tip of the choanal process
Caudal midpoint of the choanal process
Caudal deflection point between the choanal process and palatine body
Midpoint of medial margin connecting the choanal and pterygoid processes
Rostral deflection point between the pterygoid process and palatine body
Rostral midpoint of the pterygoid process
Caudal tip of the pterygoid process
Caudal midpoint of the pterygoid process
Caudal deflection point between the pterygoid process and palatine body
Lateral deflection point between the maxillary process and palatine body
Lateral midpoint of the maxillary process

**Semi-landmarks:** 15 Semi-landmarks are placed along the curve connecting landmarks 15 and 16.

## Supplementary file 2. Taxa used in geometric morphometric analysis of palatine shape

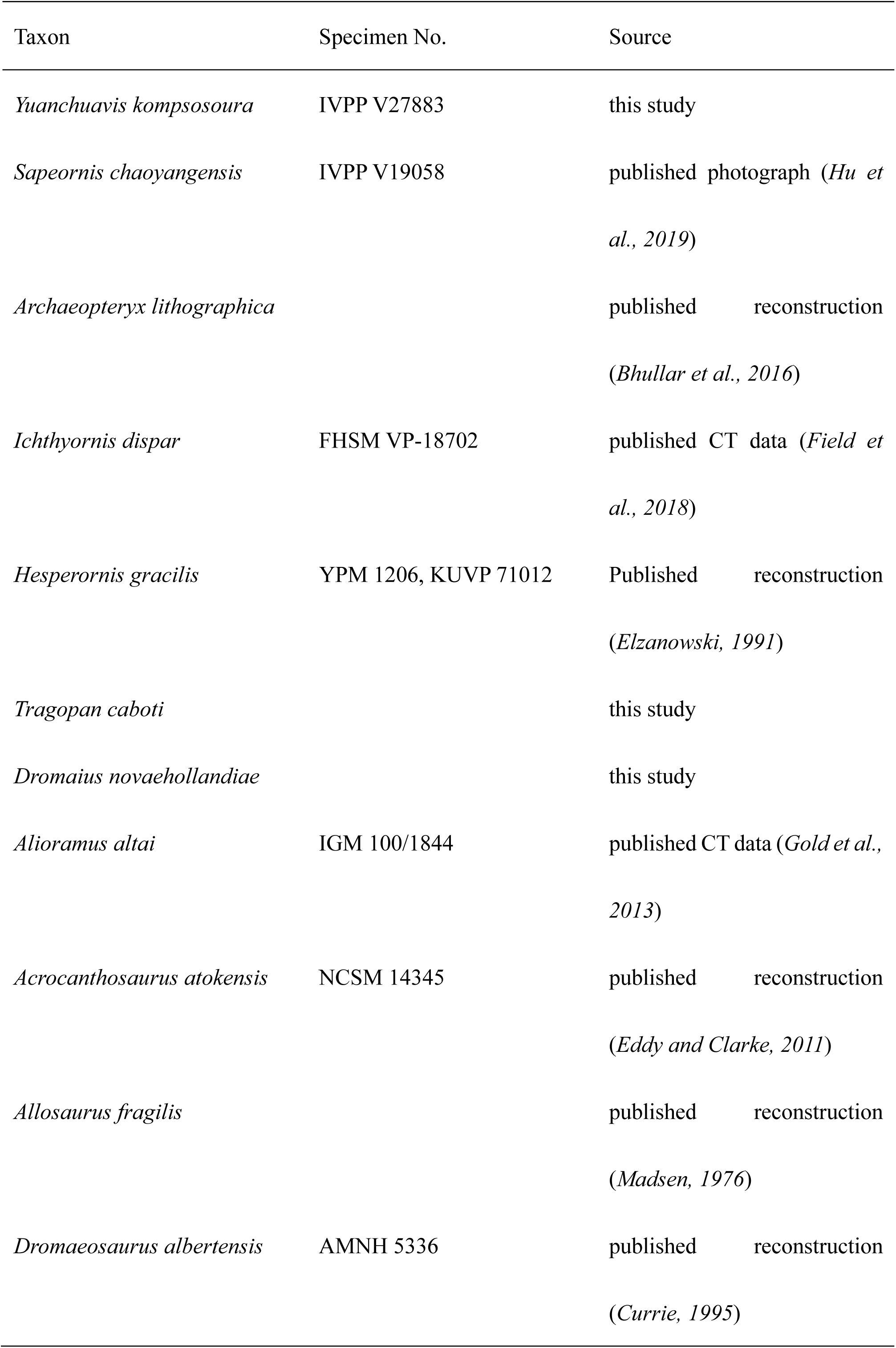
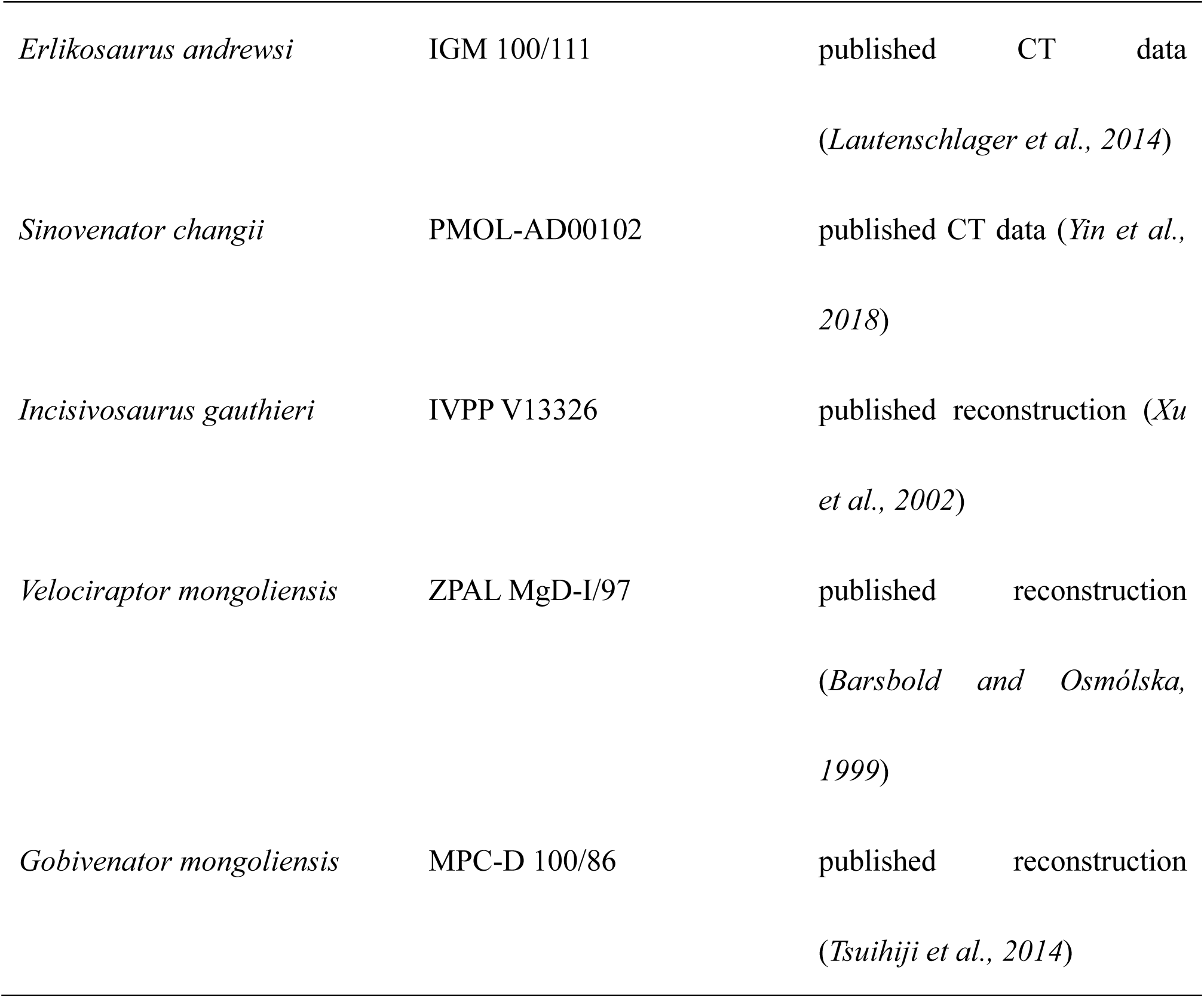

